# Comparison of single nucleotide variants identified by Illumina and Oxford Nanopore technologies in the context of a potential outbreak of Shiga Toxin Producing *Escherichia coli*

**DOI:** 10.1101/570192

**Authors:** David R Greig, Claire Jenkins, Saheer Gharbia, Timothy J Dallman

## Abstract

**Background:** We aimed to compare Illumina and Oxford Nanopore Technology (ONT) sequencing data from the two isolates of STEC O157:H7 to determine whether concordant single nucleotide variants were identified and whether inference of relatedness was consistent with the two technologies.

**Results:** For the Illumina workflow, the time from DNA extraction to availability of results, was approximately 40 hours in comparison to the ONT workflow where serotyping, Shiga toxin subtyping variant identification were available within seven hours. After optimisation of the ONT variant filtering, on average 95% of the discrepant positions between the technologies were accounted for by methylated positions found in the described 5-Methylcytosine motif sequences, CC(A/T)GG. Of the few discrepant variants (6 and 7 difference for the two isolates) identified by the two technologies, it is likely that both methodologies contain false calls.

**Conclusions:** Despite these discrepancies, Illumina and ONT sequences from the same case were placed on the same phylogenetic location against a dense reference database of STEC O157:H7 genomes sequenced using the Illumina workflow. Robust SNP typing using MinION-based variant calling is possible and we provide evidence that the two technologies can be used interchangeably to type STEC O157:H7 in a public health setting.

## Background

Shiga toxin producing *Escherichia coli* (STEC) O157:H7 is a zoonotic, foodborne pathogen defined by the presence of phage-encoded Shiga toxin genes (*stx*) [1]. Disease symptoms range from mild through to severe bloody diarrhoea, often accompanied by fever, abdominal cramps and vomiting [2]. The infection can progress to Haemolytic Uremic Syndrome (HUS), characterized by kidney failure and/or cardiac and neurological complications [3,4]. Transmission from an animal reservoir, mainly ruminants, occurs by direct contact with animals or their environment, or by the consumption of contaminated food products with reported vehicles including beef and lamb meat, dairy products, raw vegetables and salad [2,4].

STEC O157:H7 belongs to multi-locus sequence type clonal complex (CC) 11, with all but a small number of variants belonging to sequence type ST11. CC11 comprises three main lineages (I, II and I/II) and seven sub-lineages (Ia, Ib, Ic, IIa, IIb, IIc and I/II) [5]. There are two types of Shiga toxin, Stx1 and Stx2. Stx1 has four subtypes (1a-1d) and Stx2 has seven subtypes (2a-2g). Subtypes 1a, 2a, 2c, and rarely 2d, are found in STEC O157:H7. Strains harbouring *stx2a* are significantly associated with cases that develop HUS [2,6]. As well as harbouring *stx* encoding prophage, STEC O157:H7 has an additional prophage repertoire accounting for at least 20% of the chromosome.

The implementation of whole genome sequencing (WGS) data for typing STEC has improved the detection and management of outbreaks of foodborne disease [6]. Single nucleotide polymorphism (SNP) typing offers an unprecedented level of strain discrimination and can be used to quantify the genetic relatedness between groups of genomes. In general, for clonal bacteria, the fewer polymorphisms identified between pairs of strains, the less time since divergence from a common ancestor and therefore the increased likelihood that they are from the same source population. Therefore, it is paramount that variant detection for typing is accurate, highly specific and concentrated on positions of neutral evolution to ensure the correct interpretation of the sequence data within the epidemiological context of an outbreak. It has been previously shown that different bioinformatics analysis approaches for variant identification exhibit detection variability [7,8]. It is therefore important that within a particular analysis, workflow parameters to filter identified variants to achieve optimum sensitivity and specificity are appropriately optimised.

Short read sequencing platforms, such as those provided by Illumina, have been adopted by public health agencies for infectious disease surveillance worldwide [9] and have proved to be a robust and accurate method for quantifying relatedness between bacterial genomes. High-throughput Illumina sequencing although cost effective, often requires batch processing of hundreds of microbial isolates to achieve cost savings and therefore this approach offers less flexibility for urgent, small scale sequencing often required during public health emergencies [10]. In contrast, Oxford Nanopore Technologies (ONT) offers a range of rapid real-time sequencing platforms from the portable MinION to the higher throughput GridION and PromethION models, although at this time lower read accuracy compared to Illumina data suggests accurate variant calling maybe problematic.

In September 2017, Public Health England (PHE) was notified of two cases of HUS in two children admitted to the same hospital on the same night. STEC O157:H7 was isolated from the faecal specimens of both cases. In order to rapidly determine whether or not the cases were part of a related phylogenetic cluster and therefore likely to be epidemiologically linked to each other, or to any other cases in the PHE database, we sequenced both isolates using the MinION platform and integrated the ONT sequencing data with a dense reference database of Illumina sequences. We aimed to compare Illumina and ONT sequencing data from the two isolates to assess the utility of the ONT method for urgent, small scale sequencing, and to determine whether the same single nucleotide variants were identified and whether inference of relatedness was consistent with the two technologies.

## Data description

Paired-end FASTQ files were generated from the Illumina HiSeq 2500 for both samples (cases). Raw long-read data (FAST5) was generated from the MinION and basecalled using Albacore (FASTQ) in real-time. Both technologies derived FASTQ reads were trimmed and filtered (Trimmomatic, Porechop, Filtlong) before being aligned (BWA, Minimap2) to a reference genome (NC_002695.1). Variant positions were called using GATK before being imported into SnapperDB. Full processing details can be found within the methods section.

## Results

### Comparison of typing results generated by Illumina and ONT workflows

To consider the potential benefits of real-time sequencing to enhance opportunities for early outbreak detection, the timelines from DNA extraction to result generation for Illumina and ONT workflows were evaluated (Figure 1) and the relationship between yield, time and genome coverage plotted (Figure 2). For the ONT workflow, the time from DNA extraction to completion of the sequencing run was 28 hours. A total yield of 0.45 Gbases for the isolate from Case A and 0.59 Gbases for the isolate from Case B was achieved which corresponds to an equivalent coverage of the Sakai O157 STEC reference genome (5.4Mb) of 81.29X and 108.30X for isolate A and B respectively. The average PHRED quality score for all reads in Case A was 9.87 and Case B was 9.47, which is approximately 1 error every 10 bases. Base-calling and analysis was performed in real-time and serotyping, Shiga toxin subtyping and variant identification were available within six hours and twenty minutes of the 24-hour sequencing run. With respect to the Illumina sequencing workflow, the time from DNA extraction to availability of results, assuming there were no breaks in the process, was just under 40 hours (Figure 1).

**Figure 1.**
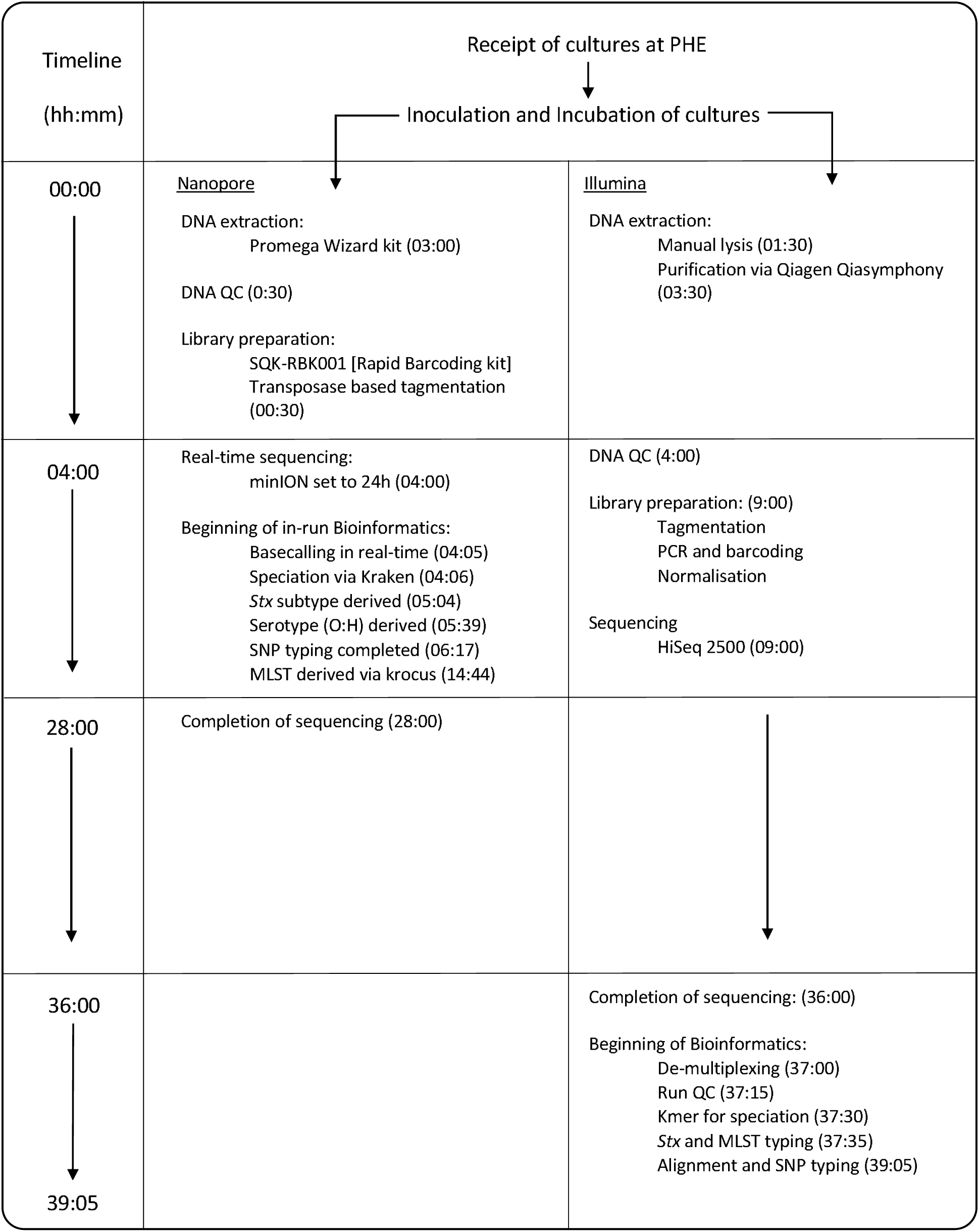
Figure showing comparative timeline from beginning DNA extraction to results generation for Oxford Nanopore and lllumina technologies. Times shown the completion of the labelled event relative to the start of the assay (hh:mm).

**Figure 2.**
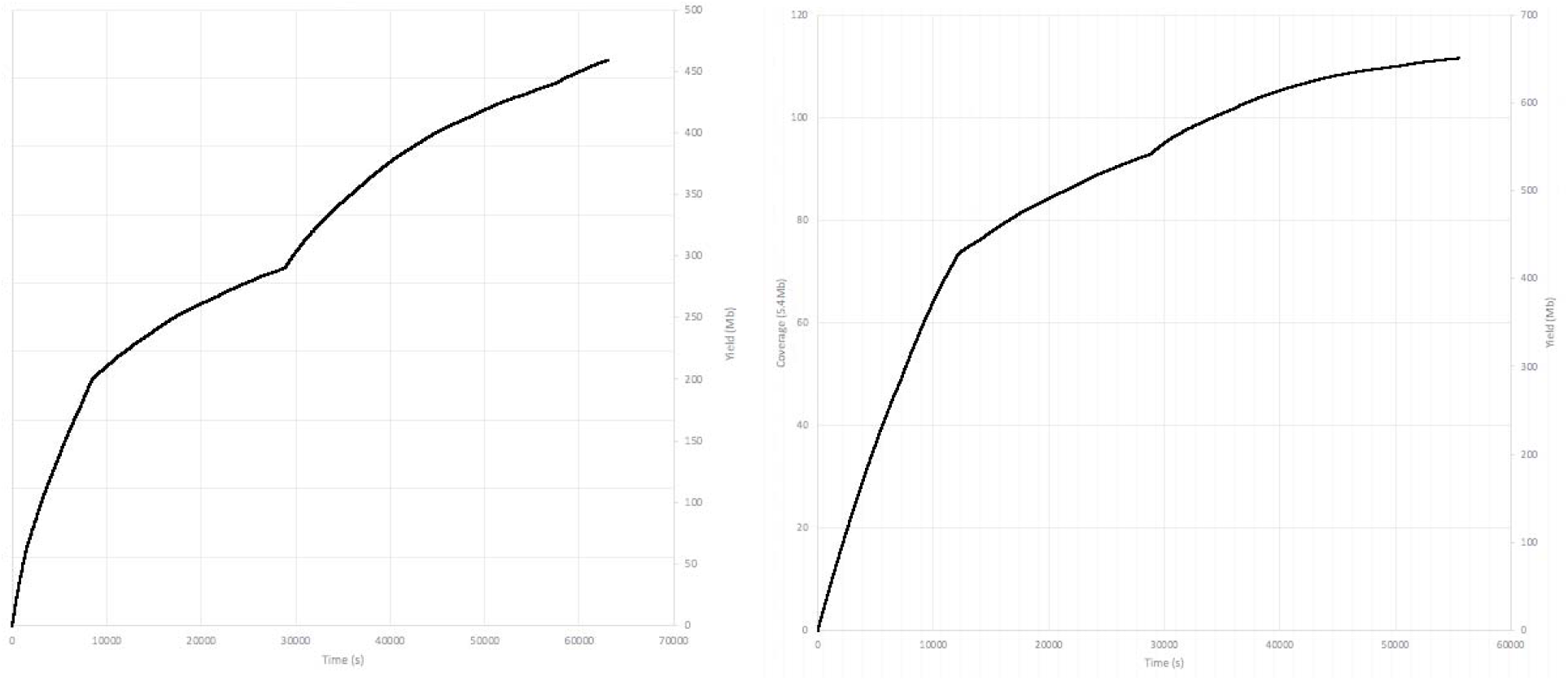
Two time/yield/coverage graphs showing production of reads in real-time and the associated cumulative mapping coverage. Case A is the graph on the left and Case B is on the right.

The species identification, serotype, MLST profile and Shiga toxin subtype results generated by both Illumina and ONT workflows were concordant with both isolates identified as *Escherichia coli* O157:H7 ST11 (12,12,8,12,15,2,2), *stx2a* and *stx2c.* During the ONT sequencing run, the bacterial species was unambiguously identified in less than one minute for both cases (Figure 1). Additionally, using Krocus, a confirmed MLST was generated for Case A at 1:54 hours and Case B at 10:39 hours into the sequencing run. This was the point at which the last read required to generate a consensus on the MLST was base-called. By 93 minutes for Case A and 41 min for Case B, it was possible to determine the *E. coli* O157:H7 serotype, and *stx*2a and *stx*2c were detected at 58 and 24 minutes into the sequencing run for Case A and Case B, respectively.

### Optimisation of ONT variant calling

To compare Illumina and ONT sequences within a standardised framework it was necessary to optimise the parameters for variant filtering within GATK2 to compensate for the lower read accuracy observed in the ONT data. Using Case B for the optimisation, base calls in the ONT data were classified as true positives (variant base detected by both methods), false positives (variant base in ONT, reference base in Illumina), true negatives (reference base in Illumina and ONT) or false negatives (variant base in Illumina, reference base in ONT). To disregard areas of the genome that the ONT reads could map to (and therefore identify variants) but were ambiguously mapped with Illumina reads, pre-filtering was performed by masking regions annotated as phage in the reference genome and those that could not be accurately self-mapped with simulated reference Illumina FASTQ reads. Figure 3 plots the precision (the proportion of true positives with respect to all positives calls) against the recall/sensitivity (the proportion of true positives identified with respect to all true positives) for an array of consensus ratio cut-offs for each of the masking strategies. Similar areas ‘under the curve’ were achieved for the different masking strategies with slightly higher precision at lower recall achieved with ‘self-masking’ (AUC – 0.71) and slightly higher recall at lower precision with explicit masking of the Sakai prophage (AUC – 0.75). The absence of a masking strategy markedly affects the precision of variant calling with ONT data, in comparison of Illumina as a gold standard (AUC – 0.30). To identify the optimum consensus cut-off for filtering ONT variants processed through GATK the F1 score was calculated at each consensus cut-off. A consensus cut-off of 0.8 maximised the precision and recall (Figure 4) irrespective of the filtering methods.

**Figure 3.**
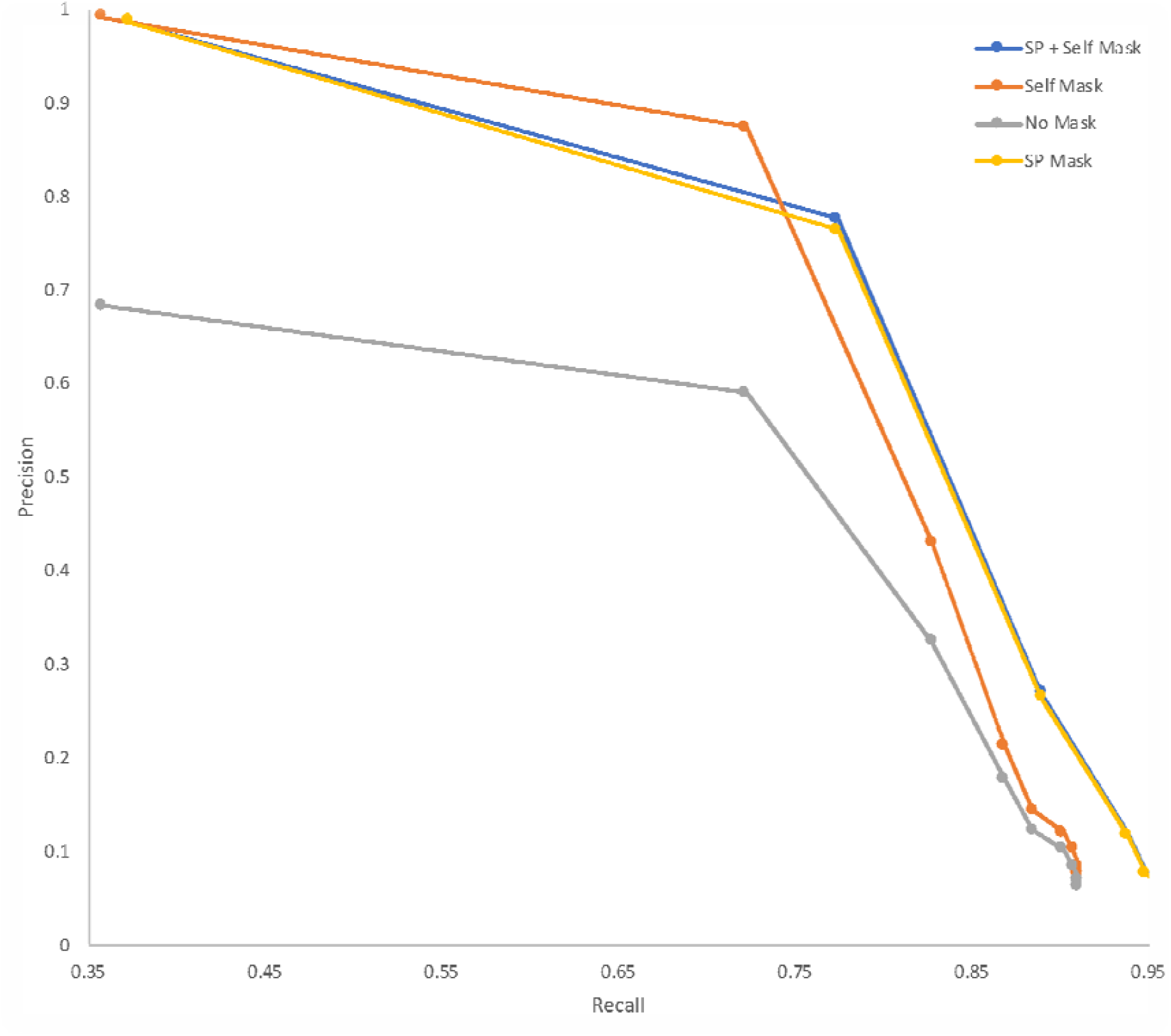
Precision Vs Recall of variant calling for an array of consensus ratio cut-offs and premasking strategies including masking positions annotated as ‘Sakai phage’ (‘SP’) and positions that are ambiguously self-mapped (‘Self’) with simulated Illumina FASTQs from the reference genome. Performed on case B.

**Figure 4.**
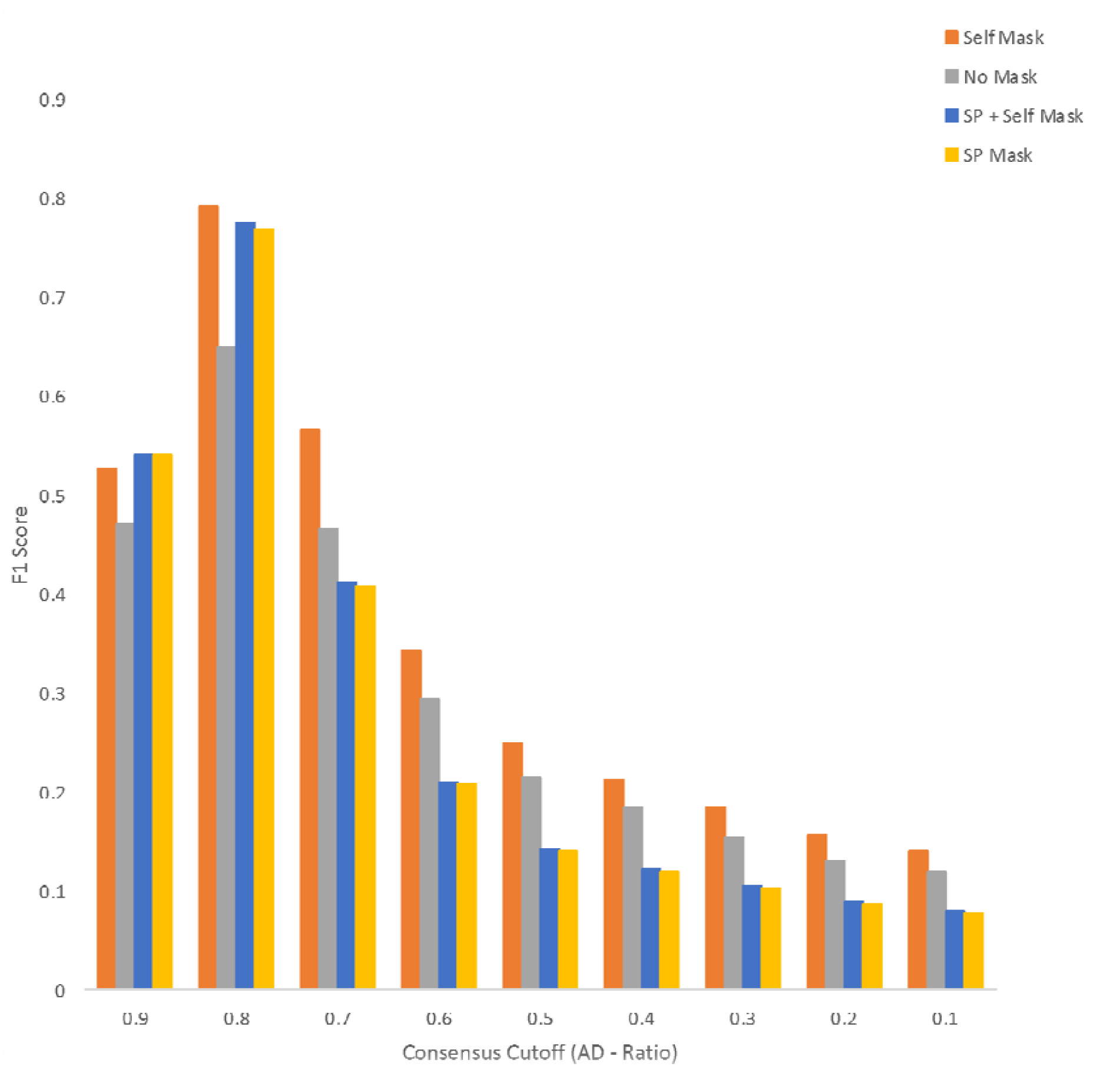
F1 Score for an array of consensus ratio cut-offs and pre-masking strategies including masking positions annotated as ‘Sakai phage’ (‘SP’) and positions that are ambiguously self-mapped (‘Self’) with simulated Illumina FASTQs from the reference genome.

### Investigation of the discrepant variants identified between the Illumina and ONT data

After optimised quality and prophage filtering there were 266 and 101 base positions for Cases A and B respectively that were discordant between the ONT and Illumina sequencing data. The majority of discrepancies were where the ONT data identified a variant not identified in the Illumina data (261/266 (98.12%) and 95/101 (94.06%) discrepant base positions for Cases A and B respectively). In contrast the Illumina data identified 5 (1.88%) discrepant base positions as variants for Case A and 6 (5.94%) for case B (Table 1) not identified by the ONT data.

**Table 1.** Table showing the breakdown of the total number of variants of each technology against the reference genome, followed by the numbers of masked variants within prophage or methylated positions.

| Variants and reason for omission. | Case A |  | Case B |  |
| --- | --- | --- | --- | --- |
|  | Illumina VCF | ONT VCF | Illumina VCF | ONT VCF |
| Total # of variants against the reference genome post quality filtering. | 2076 |  | 1424 |  |
| # of variants with masked due to location in phage | 708 |  | 531 |  |
| # of discrepant variants called between case A and B alone. | 266 |  | 101 |  |
| # of variants in each VCF. | 5 | 261 | 6 | 95 |
| # of variants with methylated positions masked. | 0 | 260 | 0 | 94 |
| Final variants. | 5 | 1 | 6 | 1 |

For both cases the most common discrepant variant were adenines classified as guanines in the ONT data with respect to the Illumina data (and reference), accounting for 68.05% (181/266) for Case A and 72.28% (73/101) for Case B. The second most common discrepancy was thymine being classified as cytosine in the ONT data accounting for 29.70% (79/266) in Case A and 22.80% (21/101) in Case B (Table 1). Of the transitions described above, 97.74% (Case A) and 93.07% (Case B) occurred when the variant was between two homopolymeric regions of multiple cytosines and guanines (Figure 5). These homopolymeric regions were similar to described DNA cytosine methylase (Dcm) binding sequences [11]. Nanopolish was subsequently used to identify likely Dcm, 5’ – cytosine – phosphate – guanine – 3’ (CpG) and DNA adenine methyltransferase (Dam) methylation sites in the ONT sequencing data and confirmed 260/266 (97.74%) and 94/101 (93.07%) discrepant variants in the ONT data were classed as methylated for Cases A and B respectively. All of which were determined to be Dcm methylation for both cases.

**Figure 5.**
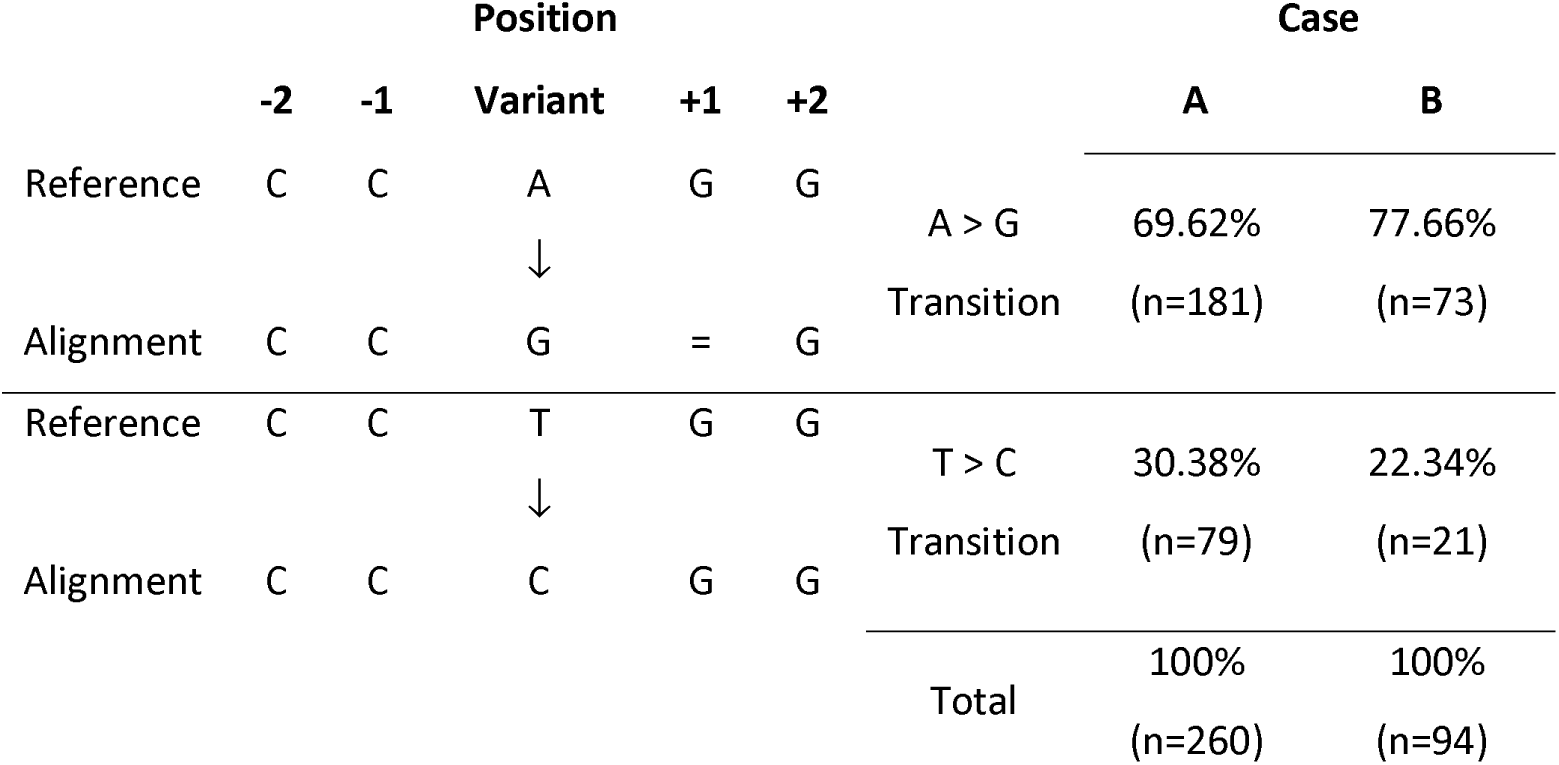
Figure showing the two most common discrepancies in the ONT optimised GATK VCFs and a breakdown of the relative proportions of these transitions compared to the total number of discrepant SNPs for both cases.

Once the methylated positions were masked from the analysis, there were a total of 6 (5 discrepant variants in Illumina and 1 ONT) and 7 (6 discrepant variants in Illumina and 1 ONT) discrepant SNPs between the ONT and Illumina data, for Cases A and B respectively (Table 2 & 3). Four discrepant Illumina variants are shared by both Case A and Case B. One shared variant was found in a non-coding region, another shared variant was found in *rhsC* encoding an RHS (rearrangement hotspot) protein defined by the presence of extended repeat regions. Two further shared variants were found in in *dadX*, an alanine racemase gene. *dadX* is a paralogue of *alr*, also annotated as an alanine racemase in the *Sakai* reference genome with significant nucleotide similarity (>75% nucleotide identity). Both intra and inter gene repeats are known to be regions of potential false positives calls with Illumina data due to miss-mapping.

**Table 2.** Table showing the final discrepant SNPs between the Illumina data and ONT data for case A. Also shown is the base as it is in the reference, the Ilumina called base and read depth at that position and the same for the ONT data. Finally, also included is the locus tag relative to the reference genome and the gene annotation.

| SNP | Position | BASE in Ref | BASE in Illum | Depth in Illum | BASE in ONT | Depth in ONT | Variant | Locus tag | Annotation |
| --- | --- | --- | --- | --- | --- | --- | --- | --- | --- |
| 1 | 270,595 | C | A | 46 | C | 141 | A | ECs0237 | rhsC |
| 2 | 379,516 | A | G | 114 | A | 100 | G | NON CODING | N |
| 3 | 1,681,338 | C | G | 59 | C | 61 | G | ECs1685 | alanine racemase 2 |
| 4 | 1,681,339 | G | C | 57 | G | 61 | C | ECs1685 | alanine racemase 2 |
| 5 | 2,636,513 | T | C | 91 | T | 69 | C | ECs2674 | hypothetical protein |
| 6 | 4,709,195 | A | A | 86 | G | 82 | G | ECs4673 | membrane-bound ATP synthase epsilon-subunit AtpC |

### Phylogenetic Analysis

Using the optimised variant calling parameters both strains clustered phylogenetically in lineage Ic within a dense reference database of STEC O157:H7 genomes (n=4475). However, the genomes were located in distinct sub-clades (Figure 6). It was, therefore, unlikely that the isolates originated from the same source, and it was concluded that Cases A and B were not epidemiologically linked. Following phylogenetic analysis of the Illumina SNP typing data (Figure 6), Case A was designated a sporadic case. However, Case B clustered with a concurrent outbreak, already under investigation, comprising three additional cases. The Illumina sequence linked to Case B was zero SNPs different from the other three cases in the cluster, whereas the ONT sequence was 7 SNPs different, when excluding the methylated positions (Table 3). Based on the ONT sequencing data alone, this discrepancy would have led to uncertainty as to whether or not the Case B was linked to the outbreak.

**Table 3.**
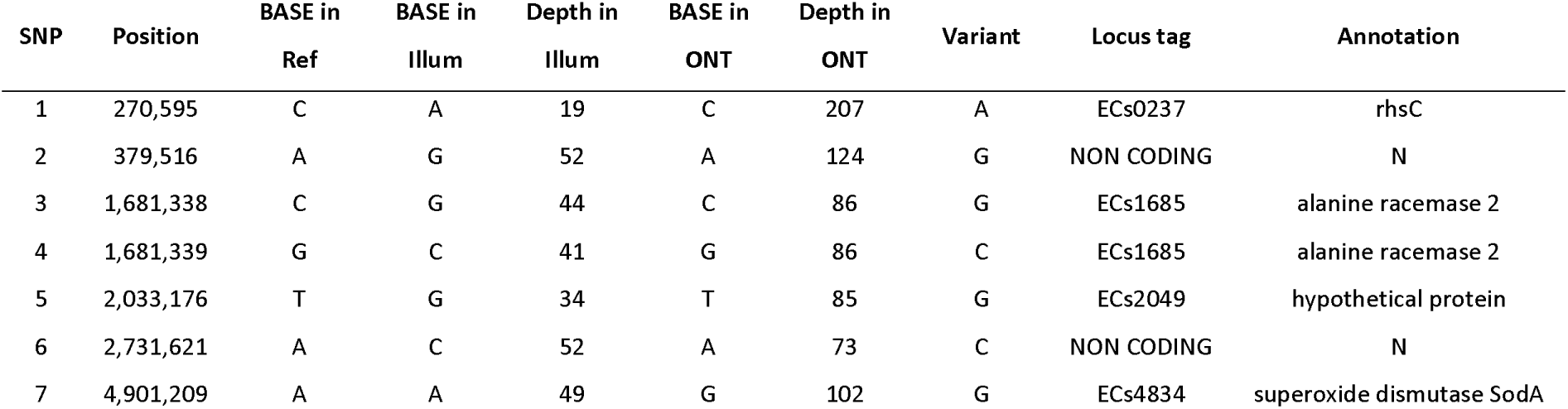
Table showing the final discrepant SNPs between the Illumina data and ONT data for case B. Also shown is the base as it is in the reference, the Ilumina called base and read depth at that position and the same for the ONT data. Finally, also included is the locus tag relative to the reference genome and the gene annotation.

**Figure 6.**
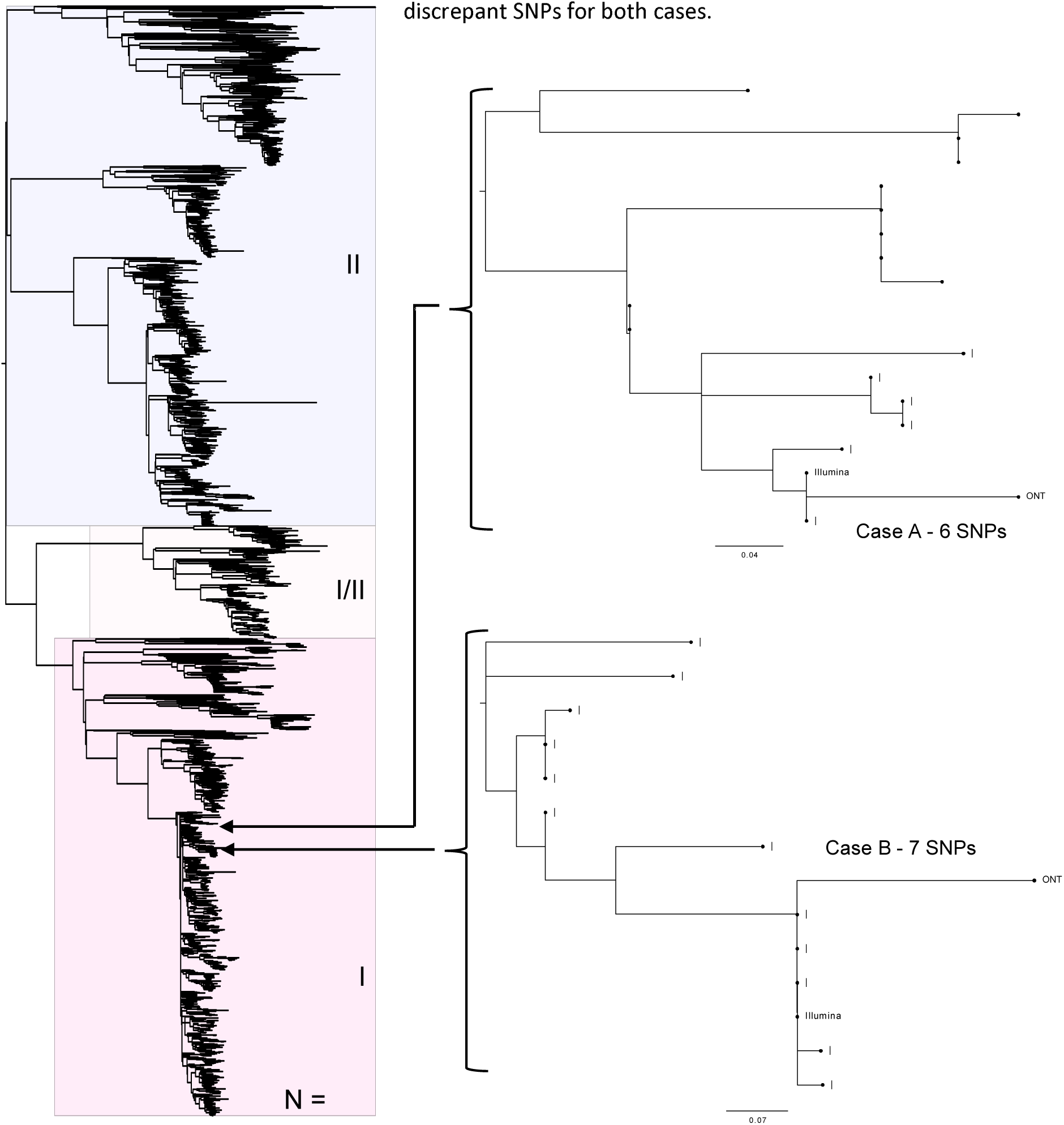
Maximum likelihood tree, of a “soft core” alignment of 4475 genomes showing the tree lineages (I, I/II and II) of STEC (Clonal Complex 11). Also showing where Oxford Nanopore and Illumina sequencing data is placed within the tree for each of the two cases. All methylated positions and prophage regions have been masked.

### Assembly Profile

The ONT-only assembly resolved to five contigs (5.73 mb) for Case A and four contigs (5.60 mb) for Case B (Supplementary Table 1). In Case A, the five contigs were determined to be a single chromosomal contig, a single plasmid contig (pO157) and the three prophage duplications. In Case B, the four contigs were determined to be a single chromosomal contig with two plasmids (one being the pO157). For Case A the assembly resolved to 25 contigs (5.51mb) with a hybrid assembly and 668 contigs (5.45 mb) with an Illumina only assembly. Case B resolved to 34 contigs (5.49 mb) with a hybrid assembly and 575 contigs (5.42 mb) with an Illumina only assembly.

Alignment of the assemblies (Supplementary Figures 1 and 2) revealed several locations within the ONT-only assembly that there were absent in the hybrid and Illumina-only assemblies. In Case A, there were 8 regions only present within the ONT-only chromosome assembly, of which 7 are related to prophage regions (Supplementary Figure 1). In case B, there were 10 chromosomal regions in the ONT-only assembly that did not align to the other assemblies. All 10 regions were associated with prophage regions (Supplementary Figures 2).

## Discussion

In this study, the two isolates sequenced using ONT were unambiguously identified as STEC O157:H7 ST 11 *stx2a*/*stx2c* in less than 15 hours and it was possible to distinguish the genetic relatedness between the isolates within 377 minutes. The WGS turn-around time from DNA extraction and library preparation, to sequencing and analysis via the Illumina workflow at PHE, is three to six days. Although this turnaround time is rapid for a service utilising batch processing on the HiSeq platforms, the sequencing approach using the MinION, whereby individual samples or small barcoded batches are loaded and results generated and analysed in real-time, has the potential to be faster and more flexible. This approach is therefore ideal for urgent, small scale sequencing, often required during public health emergencies. In this scenario, analysis of the ONT data provided evidence that the two cases were not epidemiologically linked and, although efforts were made to determine the potential source of the infection for both cases through the National Enhanced STEC Surveillance System [2], an outbreak investigation was not initiated.

A current limitation of MinION sequencing is its lower read accuracy when compared to short-read technologies [12,13,14,15,16]. This accuracy has improved as the technology has matured but still falls short of the 99% accuracy offered by short-read platforms [15]. There are a number of factors at play that contribute to the low signal to noise ratio currently inherent in the nanopore data including structural similarity of nucleotides, simultaneous influence of multiple nucleotides on the signal, the non-uniform speed at which nucleotides pass through the pore and the fact that the signal does not change within homoploymers [15]. Despite the current limitations of the technology, when mapped to references sequences in an established database of Illumina sequences, the ONT and Illumina workflow placed the sequences from the same case on the same branch in a dense reference database of STEC O157:H7 genomes sequenced using the Illumina workflow.

Although analysis of the Illumina and ONT sequencing data placed the sequences on the same branch on the phylogeny, there were SNP discrepancies between the sequences generated by the two different workflows, even after optimisation of the parameters. The vast majority of the discrepant SNPs (261/266 – 98.12% and 95/101 – 94.06 % for Cases A and B respectively) were attributed to variants identified in the ONT data and not the Illumina data. The majority of discrepancies (97.74% in Case A and 93.07% in Case B) were found in sequences that are the same as the known 5-Methylcytosine motif sequences, CC(A/T)GG [11,17] in the ONT data. Following a search of the ONT discrepant SNPs for CpG, Dam and Dcm methylation using Nanopolish, the majority (97.74% and 93.07% for case A and B respectively) of the ONT discrepant SNPs were identified in Dcm methylated regions.

As Nanopolish is detecting these methylated positions with the use of the raw FAST5 data, it is suggested that these particular discrepancies appear during the basecalling process. Albacore handles most methylation well across the three methylation models searched for by Nanopolish, for example only 94 out of 13,504 methylated positions were considered incorrect by base calling for Case B. However, for mapping based-SNP typing, this level of error in base calling means that it is not possible to accurately determine the number of SNPs, thus potentially obscuring the true phylogenetic relationship between isolates of STEC O157:H7.

The optimisation of variant filtering was performed using the Illumina data as a gold standard. However, it is possible that the alignment of the Illumina data might have generated false SNPs based on reads mapping to ambiguous regions of the genome, whereas the long reads obtained using the ONT workflow is able to resolve these ambiguous regions and call variants, or not, at these positions correctly. As the Illumina data was used as the gold standard, in this scenario SNPs produced in the Illumina data would have been classed incorrectly as false negatives in the ONT data. Discrepant variants identified in the Illumina data were attributed mainly to potentially false mapping of Illumina reads to homologous regions of the reference genome, variants which were misidentified at the same position independently in Case A and Case B. Furthermore, comparison of assemblies generated by ONT reads, Illumina reads and a hybrid approach highlights the extra genetic content accessible to ONT assemblies where variation can be quantified.

In this study an ONT sequencing workflow was used to rapidly rule out an epidemiological link between two children admitted to the same hospital on the same day with symptoms of HUS. The isolates of STEC O157:H7 from each child mapped to different clades within the same STEC O157:H7 lineage (Ic). We provide further evidence that SNP typing using MinION-based variant calling is possible when the coverage of the variation is high [15]. The error rate exhibited by ONT sequencing workflows continue to improve due to developments in the pore design, the library preparation methods, innovations in base-calling algorithms and the introduction of post-sequencing correction tools, such as Nanopolish [15,21]. Currently, both short and long read technologies are used for public health surveillance, and there is a need to integrate the outputs so that all the data can be analysed in the same way. Recently, Rang et al [15] reiterated how the scientific community can make valuable contributions to improving ONT read accuracy by systematically comparing computational strategies as highlighted in this study and elsewhere [22]. On-going up-dates to the chemistry and software tools will facilitate the robust detection of SNPs enabling ONT to compete with short read platforms, ultimately enabling the two technologies to be used interchangeably in clinical and public health settings.

## Methods

### DNA extraction, Library preparation and Illumina Sequencing

Genomic DNA was extracted from two strains of STEC O157 isolates from two HUS cases admitted to the same hospital on the same night. Using a Qiagen Qiasymphony (Qiagen, Hilden, Germany) to manufactures instructions, genomic DNA extracted and quantified using a Qubit and the BR dsDNA Assay Kit (Thermofisher Scientific, Waltham, USA) to manufactures instructions. The sequencing library was prepared by fragmenting and tagging the purified gDNA using the Nextera XT DNA Sample Preparation Kits (Illumina, Cambridge, UK) to manufactures instructions. The prepared library was loaded onto an Illumina HiSeq 2500 (Illumina, Cambridge, UK) at PHE and sequencing perfomed in rapid run mode yielding paired-end 100bp reads.

### Processing and analysis of Illumina sequence data

FASTQ reads were processed using Trimmomatic v0.27 [23] to remove bases with a PHRED score of less than 30 from the leading and trailing ends, with reads less than 50 bp after quality trimming discarded. A *k*-mer approach (https://github.com/phe-bioinformatics/kmerid) was used to confirm the species of the samples. Sequence type (ST) assignment was performed using MOST v1.0 described by [24]. *In silico* serotyping was performed by using GeneFinder, an inhouse PHE programme (Doumith, unpublished) which uses Bowtie v2.2.5 [25] and Samtools v0.1.18 [26] to align FASTQ reads to a multifasta containing the target genes (including *wzx, wzy* and *fliC*). *Stx* sub-typing was performed as described in [27]. Illumina FASTQ reads were mapped to the Sakai STEC O157 reference genome (NC_002695.1) using BWA MEM v0.7.13 [28]. Variant positions identified by GATK v2.6.5 UnifiedGenotyper [29] that passed the following parameters; >90% consensus, minimum read depth of 10, Mapping Quality (MQ) > = 30. Any variants called at positions that were within the known prophages in Sakai were masked from further analyses. The remaining variants were imported into SnapperDB v0.2.5 [30].

### DNA extraction, Library preparation and Nanopore Sequencing

Genomic DNA was extracted and purified using the Promega Wizard Genomic DNA Purification Kit (Promega, Madison, USA) with minor alterations including doubled incubation times, no vigorous mixing steps (performed by inversion) and elution into 50μl of double processed nuclease free water (Sigma-Aldrich, St. Louis, USA). DNA was quantified using a Qubit and the HS (High sensitivity) dsDNA Assay Kit (Thermofisher Scientific, Waltham, USA) to manufactures instructions. Library preparation was performed using the Rapid Barcoding Kit - SQK-RBK001 (Oxford Nanopore Technologies, Oxford, UK) with each sample’s gDNA being barcoded by transposase based tagmentation and pooled as per manufactures instructions. The prepared library was loaded on a FLO-MIN106 R9.4 flow cell (Oxford Nanopore Technologies, Oxford, UK) and sequenced using the MinION for 24 hours.

### Processing and analysis of Nanopore sequence data

Raw FAST5 files were basecalled and de-multiplexed in real-time, as reads were being generated, using Albacore v2.1 (Oxford Nanopore Technologies) into FASTQ files. Run metrics were generated using Nanoplot v1.8.1 using default parameters [31]. Reads were processed through Porechop v0.2.1 using default parameters (Wick. Unpublished) [32] to remove any barcodes and adapters used in SQK-RBK001. Samples were speciated using Kraken v0.10.4 [33]. A MLST was assigned using Krocus with the following parameters --kmer 15, --min_block_size 300 and --margin 500 [34]. S*tx* sub-typing and serotyping was determined by aligning the basecalled reads using minimap2 v2.2 [35] and Samtools v1.1 [26] to a multifasta containing the *Stx* and serotype encoding genes.

For reference based variant calling FASTQ reads were mapped to the Sakai STEC O157 reference genome (NC_002695.1) using minimap2 v2.2 [35]. VCFs were produced using GATK v2.6.5 UnifiedGenotyper [29]. Any variants called at positions that were within the known prophages in Sakai were masked from further analyses. To determine the optimum consensus cut-off for ONT variant detection the VCF was filtered with sequentially decreasing ad-ratio values at 0.1 intervals. Using the Illumina variant calls as the gold standard, F1 scores (the weighted average of precision and recall) were calculated to determine the optimal ad-ratio for processing ONT data through GATK.

### Comparison of Illumina and Nanopore discrepant SNPs

Nanopolish [21] was also used to detect methylation across the ONT data to compare to the discrepant positions. This was performed using the call-methylation function searching for three types of methylation including, the DNA adenine methyltransferase (Dam), DNA cytosine methylase (Dcm) and 5’ – cytosine – phosphate – guanine – 3’ (CpG) models. The discrepant SNPs between the Illumina and ONT for both Case A and Case B were manually visualised in Tablet v1.17.08.17 [36] in order to elucidate the reason for the discrepancy. Discordant SNPs being within a homopolymeric region were also quantified.

### Generation of phylogenetic trees

Filtered VCF files for each of the Illumina and ONT sequencing data for each sample, were incorporated, into SnapperDB v0.2.5 [30] containing variant calls from 4471 other STEC CC11 genomes generated through routine surveillance by Public Health England. SnapperDB v0.2.5 [30] was used to generate a whole genome alignment of the 4475 genomes (including both datasets for the selected strains for this study). Both methylated positions and prophage positions were masked from the alignment. The alignment was processed through Gubbins V2.0.0 [37] to account for recombination events. A maximum likelihood tree was then constructed using RAxML V8.1.17 [38].

### Assembly of ONT data

Trimmed ONT FASTQ files were assembled using Canu v1.6 [39]. Polishing of the assemblies was performed using Nanopolish v0.10.2 [21] using both the trimmed ONT FASTQs and FAST5s for each respective sample accounting form methylation using the --methylation-aware option set to dcm. Assemblies were reoriented to start at the *dnaA* gene (NC_000913) from *E. coli* K12, using the fixstart parameter in circulator v1.5.5 [40].

### Hybrid assemblies

Trimmed ONT FASTQ files were assembled using Unicycler v0.4.2 [41] with the following parameters min_fasta_length=1000, mode=normal and −1 and −2 for the incorporation of each sample’s equivalent Illumina FASTQ. Pilon v1.23 [42] was used to correct the assembly using the Illumina reads.

### Assembly of Illumina data

Illumina reads were assembled using SPAdes v3.13.0 [43] with the careful parameter activated and with kmer lengths of 21, 33, 55, 65, 77, 83 and 91.

### Annotation

Prokka v1.13 [44] with the species set to E. coli was used to annotate the final assemblies. Mauve snapshot_2015-02-25 (1) [45] using the “move contig” function was used to align each assembly to the ONT reference as they had the least number of contigs.

## Availability of supporting data

The FASTQ files for the paired read Illumina sequence data can be found on the NCBI (National Center for Biotechnology Information) Sequence Read Archive (SRA); Case A accession: SRR7184397, Case B accession - SRR6052929. The ONT FASTQ files, Case A accession – SRR7477814, Case B accession - SRR7477813. All files can be found under BioProject - PRJNA315192.

## Abbreviations

AUC: : Area under curve
BWA: : Burrows-Wheeler aligner
CC: : Clonal complex
Dam: : DNA adenine methyltransferase
Dcm: : DNA cytosine methylase
GATK: : Genome analysis toolkit
HUS: : Haemolytic Uremic Syndrome
MLST: : Multi-locus sequence type
NCBI: : National Center for Biotechnology Information
ONT: : Oxford Nanopore Technologies
PHE: : Public Health England
SNP: : Single nucleotide polymorphism
SRA: : Sequence read archive
STEC: : Shiga toxin-producing *Escherichia coli*
VCF: : Variant call format
WGS: : Whole genome sequencing

## Author contributions

CJ and TJD conceptualised the project. CJ and DRG performed DNA extractions. DRG performed library preparation and Nanopore sequencing. TJD and DRG processed Illumina sequence data. DRG processed all ONT data. TJD performed ONT optimisation. DRG performed methylation analysis. TJD and DRG performed Illumina and ONT data comparison. CJ wrote the original draft. DRG, CJ, SG and TJD performed manuscript editing.

## Acknowledgements

We would like to thank Oxford Nanopore Technologies for funding this research. In particular we would like to thank Leila Luheshi and Divya Mirrington for their support and scientific assistance. We would also like to thank the frontline NHS Laboratories for submitting the samples used in this study to the Gastrointestinal Bacteria Reference Unit at Public Health England.

We would like to acknowledge Dr Andrew Page of the Quadram Institute for critically reviewing the manuscript.

## Supplementary Tables

**Table 1.** Table showing the number of contigs generated and size of assembly for each assembly method for both cases.

| Case | # of contigs in ONT-only assembly (size bp) | # of contigs in hybrid assembly (size bp) | # of contigs in Illumina-only assembly (size bp) |
| --- | --- | --- | --- |
| A | 5 (5,725,666 bp) | 25 (5,506,670 bp) | 668 (5,449,735 bp) |
| B | 4 (5,620,611 bp) | 34 (5,491,608 bp) | 575 (5,424,436 bp) |

## Supplementary Figures

**Supplementary figure 1.**
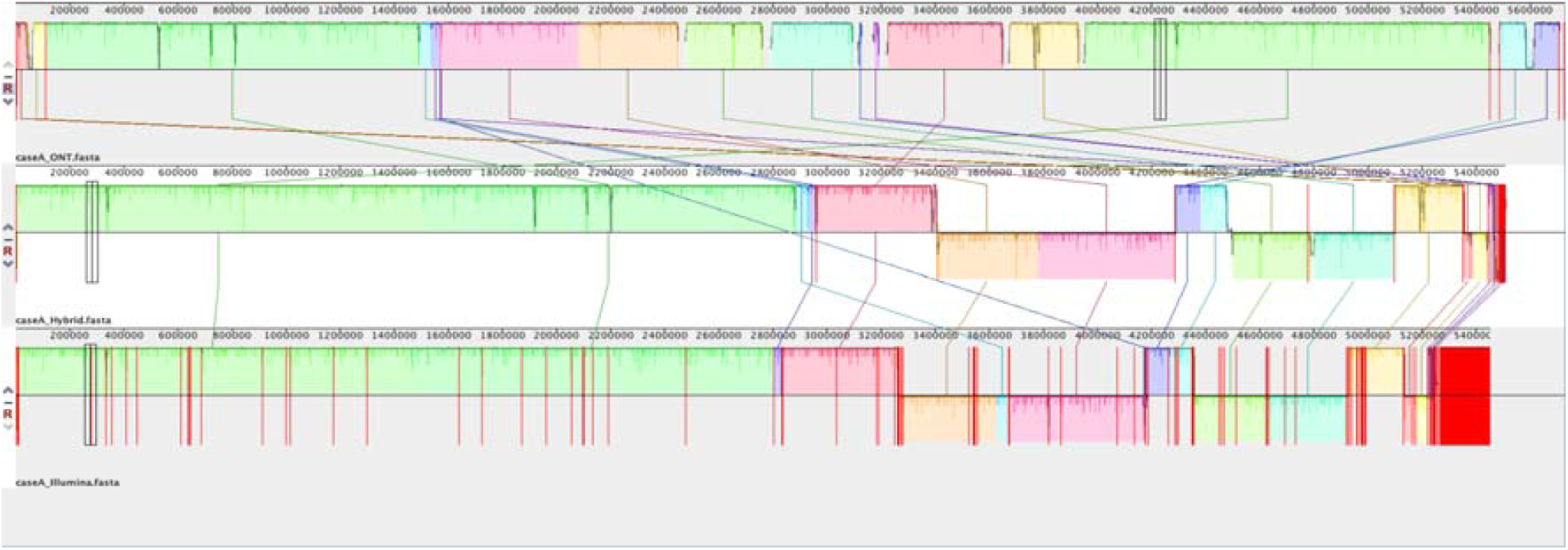
Mauve alignment showing regions of similarity between the ONT-only, hybrid and Illumina-only assemblies (order descending) for Case A. Also showing the chromosomal regions in the ONT-only assembly that did not match the other assemblies (red arrows).

**Supplementary figure 2.**
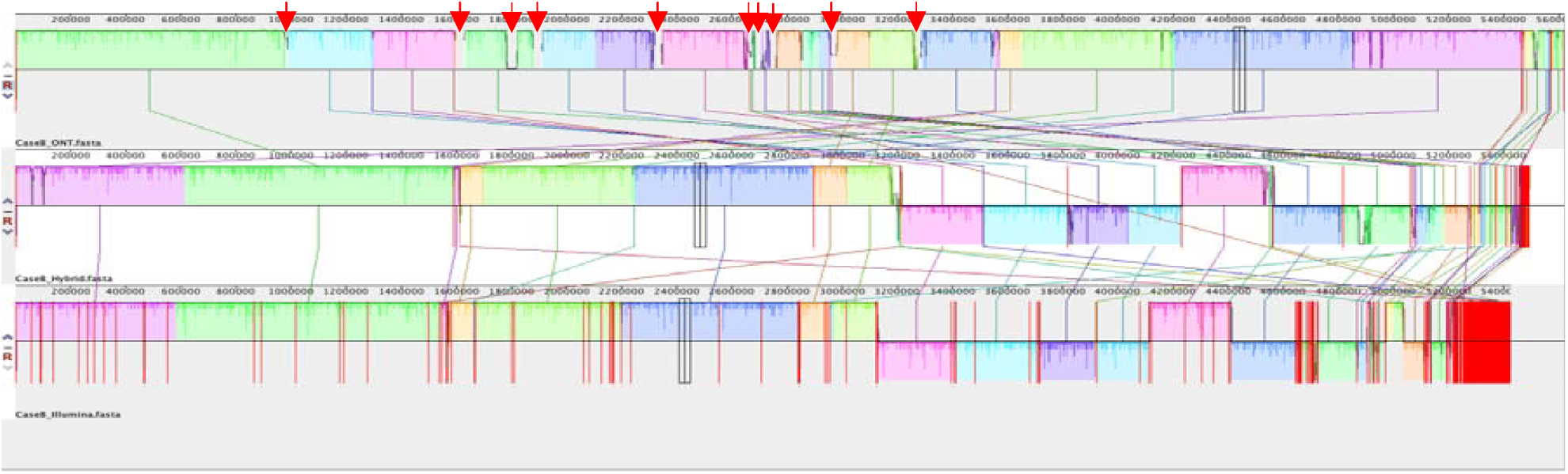
Mauve alignment showing regions of similarity between the ONT-only, hybrid and Illumina-only assemblies (order descending) for Case B. Also showing the chromosomal regions in the ONT-only assembly that did not match the other assemblies (red arrows).

